# Rapamycin induced hyperglycemia is associated with exacerbated age-related osteoarthritis

**DOI:** 10.1101/2021.05.21.445179

**Authors:** Dennis M. Minton, Christian J. Elliehausen, Martin A. Javors, Kelly S. Santangello, Adam R. Konopka

## Abstract

**Background:** The objective of this study was to determine if mechanistic target of rapamycin (mTOR) inhibition with or without AMP-activated protein kinase (AMPK) activation can protect against primary, age-related OA.

**Design:** Dunkin-Hartley guinea pigs develop mild primary OA pathology by 5-months of age that progresses to moderate OA by 8-months of age. At 5-months, guinea pigs sacrificed as young control (n=3) or were fed either a control diet (n=8), a diet enriched with the mTOR-inhibitor rapamycin (Rap, 14ppm, n=8), or Rap with the AMPK-activator metformin (Rap+Met, 1000ppm, n=8) for 12 weeks. Knee joints were evaluated by OARSI scoring, micro-computed tomography, and immunohistochemistry. Glenohumeral articular cartilage was collected for western blotting.

**Results:** Rap and Rap+Met treated guinea pigs displayed lower body weight than control. Rap and Rap+Met inhibited articular cartilage mTORC1 but not mTORC2 signaling. Rap+Met, but not Rap alone, stimulated AMPK. Despite lower body weight and articular cartilage mTORC1 inhibition, Rap and Rap+Met treated guinea pigs had greater OA severity in the medial tibial plateau due to articular cartilage structural damage and/or proteoglycan loss. Rap and Rap+Met increased plasma glucose compared to control. Plasma glucose concentration was positively correlated with proteoglycan loss, suggesting hyperglycemic stress may have contributed to worsened OA.

**Conclusions:** This is the first study to show that Rap induced increase in plasma glucose was associated with greater OA severity. Further, articular cartilage mTORC1 inhibition and bodyweight reduction by dietary Rap and Rap+Met did not protect against primary OA during the prevailing hyperglycemia.

## Background

Primary, age-related osteoarthritis (OA) is estimated to account for as many as 90% of all knee OA cases in humans (1). However, preclinical research commonly relies on experimental models of secondary OA. Although primary and secondary OA share similar pathological outcomes, there is a growing body of evidence to suggest they are driven by distinct mechanisms. Retrospective analysis of differentially expressed genes from separate cohorts of primary and secondary OA patients relative to their healthy controls found that only 10% of differentially upregulated and 35% of differentially downregulated genes in OA vs non-OA samples are conserved between primary and secondary OA (2,3). Therefore, 65-90% of differentially expressed genes may be unique to primary versus secondary OA. Additionally, transgenic animal models have revealed that several genes are differentially involved in the progression of primary and secondary OA (4–9). For example, deletion of *Panx3* protects against secondary OA yet dramatically worsens primary OA (4), and deletion of *JNK1/2* accelerates the development of primary OA while having no effect on secondary OA progression (9). Together, these studies reinforce that unique mechanisms underpin these two forms of OA.

Age is one of the greatest risk factors for nearly every chronic disease, including primary OA. Two evolutionarily conserved kinases, mechanistic target of rapamycin (mTOR) and AMP-activated protein kinase (AMPK), are energy sensing pathways similarly dysregulated during aging and OA (10–13). The mTOR inhibitor rapamycin (Rap) can extend lifespan in mice and delay the onset of several age-related morbidities (12,14). The anti-diabetic drug metformin (Met) can activate AMPK and, when added to Rap, extends lifespan to a greater extent than historical cohorts of mice treated with Met or Rap alone (15). Additionally, Met is the first drug being tested to slow age-related multi-morbidity in humans (16). While the prospect of lifespan extension is tantalizing, extending lifespan without delaying the onset or slowing the progression of the most debilitating age-associated conditions could be viewed as detrimental. Therefore, it is imperative to understand if purported lifespan-extending therapies that target the fundamental biology of aging are also capable of delaying the onset of chronic diseases, such as primary OA.

mTOR exists as complex I (mTORC1) and complex II (mTORC2). mTORC1 regulates cellular proliferation, protein synthesis, senescence, and survival while mTORC2 functions downstream of insulin signaling on substrates such as PI3K-Akt (12). In articular cartilage, mTORC1 activity increases with age and is sufficient to induce OA in young male mice (10). In non-articular tissues, acute or intermittent Rap selectively inhibits mTORC1 while chronic Rap administration for durations greater than 14 days also inhibits mTORC2 activity (17). Cartilage-specific deletion of mTOR and systemic or intra-articular injections of Rap and the mTORC1/2 inhibitor Torin 1 lower secondary OA in young-male mice and rabbits (18–21). While these findings support mTOR-based therapeutics for OA, the completed studies were exclusively in injury-induced models of OA and have not been investigated in primary, age-related OA.

Recently, it has been proposed that the positive effects of mTOR inhibition on OA pathology may be diminished by feedback activation of PI3K and has raised questions about the need for a dual treatment strategy that inhibits both mTOR and upstream PI3K signaling (22,23). In addition to activating AMPK, Met has pleotropic effects including inhibition of PI3K signaling in rheumatoid arthritis fibroblast-like synoviocytes (24). Moreover, Met and other AMPK-activators have chondroprotective effects against inflammatory-induced protease expression *in vitro* (25,26) and protect against injury-induced OA in young male mice and rhesus monkeys (27). Treatment with Met is also is associated with a lower rate of medial tibiofemoral cartilage volume loss and risk of total knee replacement in obese patients (28). However, Met as an adjuvant therapy to Rap has not been investigated in primary OA.

The Dunkin-Hartley guinea pig is a well-characterized outbred model of primary OA. The progression of OA in guinea pigs is related to bodyweight (29) and shares a similar age-related and spatial progression to humans (30). Mild OA pathology develops by 5 months in guinea pigs that progresses to moderate OA by 8-9 months of age (30–32). Therefore, at 5 months of age we treated guinea pigs with lifespan-extending doses of Rap or a combination of Rap+Met for 12 weeks to slow the progression from mild to moderate OA. This study is the first to evaluate if lifespan extending treatments can modify primary OA, the most prevalent form of OA observed in older adults.

## Methods

### Animal Use

All tissues were collected at the University of Illinois Urbana-Champaign and approved by the Institutional Animal Care and Use Committee. Data collection and analysis were completed at University of Wisconsin-Madison and William S. Middleton Memorial Veterans Hospital. Because male Dunkin-Hartley guinea pigs develop more severe OA pathology than female (33), we used male animals to maximize the potential for the interventions to slow the progression of OA. Therefore, similar to previous work (34), male Dunkin-Hartley guinea pigs (Charles River) were singly housed in clear plastic, flat bottomed cages (Thoren, Model #6) with bedding. Guinea pigs were single housed to measure food consumption. 12-hour light/dark cycles were used beginning at 0600. Guinea pigs acclimated for 2-3 weeks and were provided standard chow diet (Evigo 2040) fortified with vitamin C (1050 ppm) and Vitamin D (1.5 IU/kg) and water ad libitum until 5 months of age. Guinea pigs were then sacrificed to serve as young control (n=3), randomized to continue the standard diet (n=8), or receive standard diets enriched with encapsulated rapamycin (14 ppm, n=8) or the combination of encapsulated rapamycin and metformin (14 ppm, 1000 ppm, n=8) for 12 weeks. Guinea pigs were randomized to match bodyweight between groups prior to beginning treatment. Diets were enriched with microencapsulated rapamycin (Rapamycin holdings) and/or metformin (AK Scientific, I506) at concentrations previously shown to extend lifespan in mice (14,15,35). Food consumption was recorded on Monday, Wednesday, and Friday between 8 and 9 AM, and body weight was recorded before feeding on Monday. Guinea pigs treated with Rap or Rap+Met diet had ad libitum access to food. Dietary Rap treatment has been shown to significantly reduce bodyweight in mice (36,37). Therefore, we matched food consumption in the control group to the Rap diets to minimize the influence of food intake on dependent variables. One guinea pig in the Rap+Met group was euthanized early due to a wound on the gums which led to suppressed appetite and infection. Tissues from this animal were not collected for analysis. It could not be determined if this was due to a laceration or an oral ulcer, the latter of which is a known side effect of mTOR inhibitors (38).

### Tissue Collection

Two animals were sacrificed daily between 7 and 10 AM. Food and water were removed from the cages 2-4 hours before euthanasia. Animals were anesthetized in a chamber containing 5% isoflurane gas in oxygen and maintained using a face mask with 1.5-3% isoflurane. Blood was collected by cardiac venipuncture followed by excision of the heart. The right hind limb was removed at the coxofemoral joint, fixed in 10% neutral buffered formalin (NBF) for 48 hours, and transferred to 70% ethanol until processed for histology. Glenohumeral cartilage was collected, snap frozen in liquid nitrogen, and stored at −80C for further analysis. Because testicular atrophy has been observed following Rap treatment (39), the left testicle was preserved in 10% NBF and weighed. Although tissues are commonly weighed before fixation, previous work demonstrates that fixation negligibly effects testicle weight in similarly sized rodents (40).

### Analysis of Experimental Diets and Blood

Samples of diets enriched with Rap, Met or the combination of Rap+Met, and aliquots of whole blood (n=4 per group) were sent to the Bioanalytical Pharmacology Core at the San Antonio Nathan Shock Center to confirm drug concentrations in the diet and in circulation. Analysis was performed using tandem HPLC-MS as described previously (14,41,42). Frozen aliquots of plasma were thawed to measure glucose and lactate concentrations using the YSI Biochemistry Analyzer (YSI 2900).

### Micro Computed Tomography (µCT)

Right hind limbs from half of each treatment group (n=4 per group) were scanned using a Rigaku CT Lab GX130 at 120 μA and 110 kV for 14 minutes, achieving a pixel size of 49 μm. Scans were first processed in Amira 6.7 (ThermoFisher) where epicondylar width was measured and a series of dilation, erosion, filling, and image subtraction functions were used to isolate trabecular and cortical bone as described previously (43). Scans were then resliced 4 times along axes perpendicular to medial and lateral tibial and femoral articular surfaces and binarized using identical thresholds. NIH ImageJ software and BoneJ plugin were used to quantify thickness, spacing, and volume fraction measurements. Cortical thickness was measured by placing polygonal regions of interest (ROI) in resliced scans to encompass the articular surfaces in each joint compartment. Trabecular thickness, spacing, and bone volume fraction were measured by placing transverse ROIs (2.4×2.4×1mm) in the trabecular bone of each joint compartment.

### Histology

Knee joints were decalcified in a 5% ethylenediaminetetraacetic acid, changed every 2-3 days for 6 weeks. Joints were then cut in a coronal plane along the medial collateral ligament, paraffin embedded and sectioned at 5um increments for Toluidine Blue staining and immunohistochemistry (IHC). Slides were scanned using the Hamamatsu NanoZoomer Digital Pathology System, providing 460nm resolution. Scan focus points were set manually along the articular cartilage. Imaged slides were then scored by two blinded reviewers for OA severity following OARSI Modified Mankin guidelines as described (32). Briefly, toluidine blue stained histology slides were assigned scores for severity of articular cartilage structural damage (0-8), proteoglycan loss as assessed by absence of toluidine blue staining (0-6), disruption of chondrocyte cellularity (0-3), and tidemark integrity (0-1), with a total possible score of 18 per joint compartment (Total OARSI Score). One guinea pig each from the Rap and Met groups were unable to be analyzed due to off-axis transection before embedding. One control animal was a statistical outlier as detected by Grubb’s test and was excluded from the study. Therefore, n=7 per group were used for histopathological analysis.

### Immunohistochemistry

Antigen retrieval was performed in 10mM sodium citrate for 7 hours at 60C. Endogenous peroxidase activity was quenched using 3% H_2_O_2_ for 15min before blocking in 5% normal goat serum diluted in TBST for 1 hour at RT. Slides were incubated overnight in 200-300 uL of either p-RPS6 (1:200 dilution; Cell Signaling, 4858) or a rabbit IgG isotype control (Cell Signaling, 3900) diluted to match primary antibody concentration. Primary antibodies against p-Akt Ser473 (1:100 dilution; 4060) and p-AMPK Thr172 (1:200 dilution; 50081) from Cell Signaling were attempted, but reactivity was not seen in guinea pig articular cartilage. 150-200uL of goat anti-rabbit secondary antibody (Cell Signaling, 8114) was added for 1 hour at room temperature followed by exposure in 3,3’-diaminobenzadine (DAB; Cell Signaling, 8059) for 10 minutes. Slides were then counterstained using hematoxylin, dehydrated, and cleared through graded ethanol and xylene, coverslipped using Permount (Electron Microscopy Sciences), and viewed and imaged under a brightfield microscope. No DAB staining was seen following incubation with the IgG control or secondary antibody alone, confirming specificity of the primary antibody. For quantification, ROIs were placed to encompass areas of staining in the medial tibial articular cartilage, and cells were counted to determine the percent-positive cells. For intensity-based quantification, a color deconvolution for DAB staining was applied in ImageJ, and mean integrated intensity was quantified by averaging two p-RPS6 replicates and subtracting background staining of IgG controls.

### Western Blot

Cartilage was removed from the glenohumeral joint using a scalpel and placed in reinforced Eppendorf tubes containing 500 mg of ceramic beads (Fisher, 15-340-160) and 200 μL of RIPA buffer with protease and phosphatase inhibitors (Sigma, 5892970001), and homogenized by 2, 30-second cycles at 6 m/s in the Omni BeadRuptor. Homogenate was transferred to microcentrifuge tubes and spun at 10,000g for 10 min at 4C. Supernatants were diluted to equal concentration following a BCA assay. Samples were prepared in reducing conditions with β-mercaptoethanol in 4x Laemmli Sample Buffer (BioRad, 1610747) and heated at 95C for 5 minutes. 10 μg of protein was separated on 4-15% TGX precast gels (BioRad, 4561083) and transferred to PVDF membranes (BioRad, 1620177). Membranes were blocked in TBST with 5% bovine serum albumin (Sigma, A9647) for 1 hour at RT and incubated overnight at 4C in primary antibodies against p-RPS6 Ser235/236 (4858), RPS6 (2217) p-Akt Ser473 (4060), Akt (4685), P-AMPK Thr172 (50081), AMPK (2532), and LC3B (3868) from Cell Signaling and ADAMTS5 (ab41037), MMP-13 (ab39012), and b-Actin (ab8226) from Abcam. HRP-conjugated anti-Rabbit (Cell Signaling) or anti-Mouse (Abcam) secondary antibodies were diluted 1:5,000 for all proteins except b-Actin (1:10,000 dilution). All membranes were imaged using a UVP BioSpectrum 500 (UVP) following 5-minute incubation in a 2:1 combination of SuperSignal Pico (Fisher, 34577) and Femto (Fisher, PI34095) chemiluminescent substrates except b-Actin which received Pico alone. Densitometric analysis was performed using VisionWorks (Analytikjena). Phosphorylated proteins are expressed relative to their total protein and other targets are expressed relative to b-Actin.

### Statistical Analysis

Previous work demonstrated that a sample size of n=6 is adequately powered to detect changes between groups in guinea pigs (34). Therefore, we a priori determined our sample size (n=7-8 per group) to be appropriate to detect differences between treatment groups. All data were subjected to normality testing via the Shapiro-Wilk test. Comparisons of normally distributed data were performed using two-way unpaired t-tests or one-way ANOVA followed by Holm-Sidak’s multiple comparison test. Data with non-Gaussian distribution were compared using non-parametric Mann-Whitney tests or the Kruskal-Wallis test followed by Dunn’s multiple comparisons test. A two-way repeated measures ANOVA (time x treatment) was performed to determine differences in food consumption and body weight. Upon a significant interaction, Holm-Sidak’s multiple comparisons test was used. Because we were interested in determining if treatments impacted the trajectory of OA pathogenesis compared to aged controls, differences in all other variables besides plasma glucose were made using one-way ANOVA comparing treatment groups to 8-month controls. Due to previous reports that Met can rescue the hyperglycemic effects of Rap (37), comparisons were made between all groups for plasma glucose. Pearson’s R was used to determine correlation between variables. P-values <0.05 were considered statistically significant. Data are presented as scatter plots with mean or mean ± standard deviation (SD).

## Results

### Influence of rapamycin and rapamycin+metformin on guinea pig physical and metabolic characteristics

Figure 1A shows the average daily food consumption per week of standard diet or standard diet enriched with Rap or Rap+Met. The average daily intakes of Rap and Met based on food consumption and dietary concentration are reported in Table 1. Compared to control, there was decreased food consumption in guinea pigs receiving Rap+Met during week 2 (P=0.04). There were no significant differences between treatments. Despite largely matching food intake, there was a significant effect for treatment (P=0.004) and an interaction between time and treatment (P<0.0001) on bodyweight. Rap+Met (P=0.01) and Rap-treated guinea pigs (P=0.02) were smaller than control starting at week 3 and week 4, respectively, until the end of the study (Figure 1B). At sacrifice, Rap (P=0.002) and Rap+Met-treated guinea pigs (P=0.001) were 15% and 22% smaller than control.

**Table 1:** Average consumption of rapamycin and metformin. Using the concentration of rapamycin and metformin from the diet analysis, the average doses were calculated for each group. N=7-8 per group. Data are presented as mean ± SD.

|  | Experimental Diet |  |
| --- | --- | --- |
|  | Rapamycin | Rapamycin+Metformin |
| Rapamycin consumed (mg/kg/day) | 0.72 ± 0.09 | 0.68 ± 0.08 |
| Metformin consumed (mg/kg/day) | - | 45 ± 5.6 |

**Figure 1:**
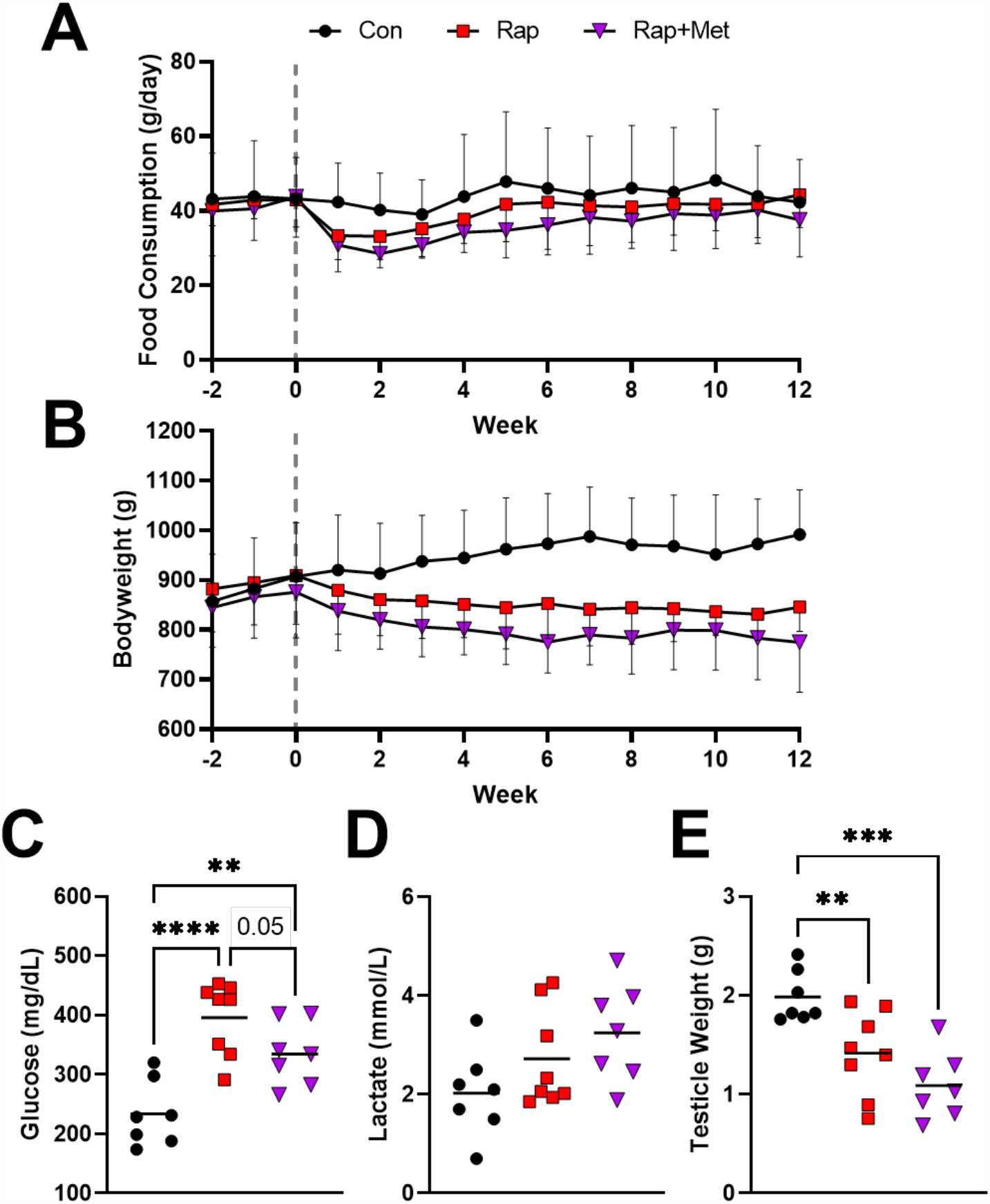
Characterization of animals on experimental diets. Food consumption (A) and bodyweight (B) of guinea pigs were recorded for the duration of the study (data presented as mean with shaded bands representing SD). Plasma glucose (C), lactate (D), and testicle weight (E) are shown. **P<0.01 vs Con, ***P<0.001 vs Con, ****P<0.0001 vs Con.

Treatment with Rap (396±61 mg/dL; P<0.0001) and Rap+Met (334±53 mg/dL; P=0.007) increased plasma glucose compared to control (234±55 mg/dL), and the addition of Met to Rap decreased plasma glucose compared to Rap alone (P=0.05; Figure 1C). Lactate concentration trended to be elevated by 66% in Rap+Met-treated guinea pigs, only (P=0.07; Figure 1D). Testicle weight in guinea pigs receiving Rap (P=0.006) and Rap+Met (P=0.0003) were 27% and 44% lower than control, respectively, suggesting gonadal atrophy (Figure 1E). We analyzed blood for the circulating Rap and Met concentrations ∼3-hours after food had been removed from the cage (Table 2). This timing aligns with a measurement of peak circulating Rap and Met. We saw that experimental diets were sufficient to increase Rap and Met concentrations in the blood, and that Rap values were not different when providing diets individually or in combination. There was no Rap or Met detected in circulation in control animals.

**Table 2:**
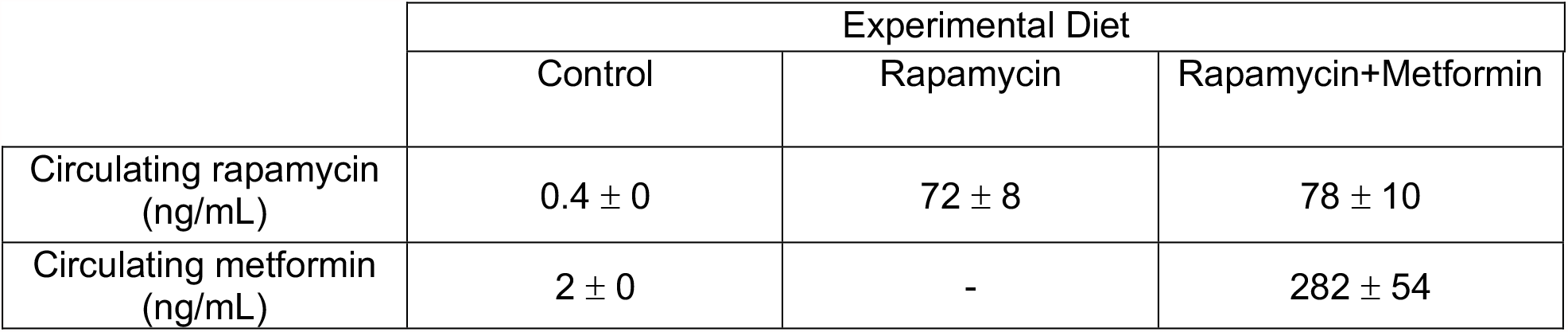
Concentrations of rapamycin and metformin in circulation. Whole blood was collected ∼3 hours after food had been removed from the cages of guinea pigs and was analyzed for rapamycin and metformin concentration by tandem HPLC/MS. N=4 per group. Data are presented as mean with ± SD.

### Rapamycin and rapamycin+metformin treatment exacerbated the age-related progression of OA

Consistent with the age-related progression of mild to moderate OA in guinea pigs, we observed an increase in medial tibial total OARSI score from 5 to 8 months (P=0.03; Figure S1A-B). Surprisingly, Rap and Rap+Met treatment resulted in a ∼2-fold increase in total OARSI score in the medial tibial plateau compared to 8 month old, age-matched control (P=0.02 for both Rap and Rap+Met; Figure 2B). This was driven by increased scores for articular cartilage structure (P=0.02 for Rap, P=0.11 for Rap+Met; Figure 2C) and proteoglycan loss (P=0.02 for Rap and Rap+Met; Figure 2D). There was no significant effect of Rap or Rap+Met on the OARSI score for the lateral tibia or medial or lateral femur (Figure S1C).

**Figure 2:**
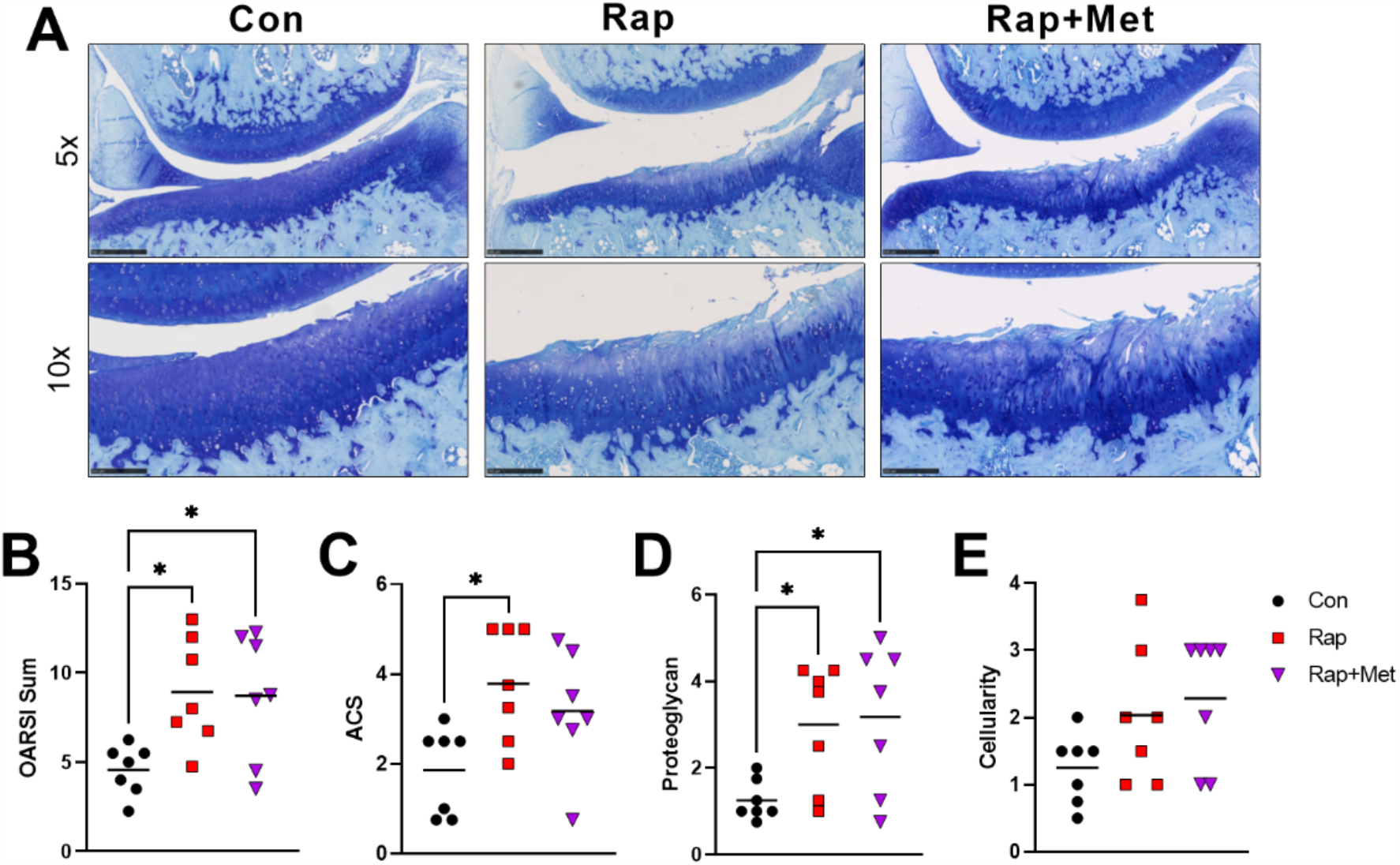
Rapamycin and rapamycin plus metformin worsened primary OA. Representative images of histology from the medial tibia are shown for each group (A; scale bars are 0.5mm and 0.25mm in 5x and 10x images, respectively). Histological images were graded for total OARSI score (B; n=7 per group). The individual scores for articular cartilage structure (C), proteoglycan loss (D) and cellularity (E) are also shown. *P<0.05 vs Con.

### OA pathology was correlated to plasma glucose, bodyweight, and testicle weight

Because Rap and Rap+Met treated guinea pigs displayed several side effects of Rap, including increased plasma glucose, testicular atrophy, decreased bodyweight, and worsened OA pathology, we evaluated the relationship between these variables and measures of OA severity across all guinea pigs. Plasma glucose was positively correlated to proteoglycan loss (R^2^=0.19; P=0.04; Figure 3A), and total OARSI score was negatively correlated with both bodyweight (R^2^=0.19; P=0.04; Figure 3B) and testicle weight (R^2^=0.20; P=0.04; Figure 3C). However, because testicle weight and bodyweight were also related (data not shown), the individual contribution of these variables cannot be resolved.

**Figure 3:**
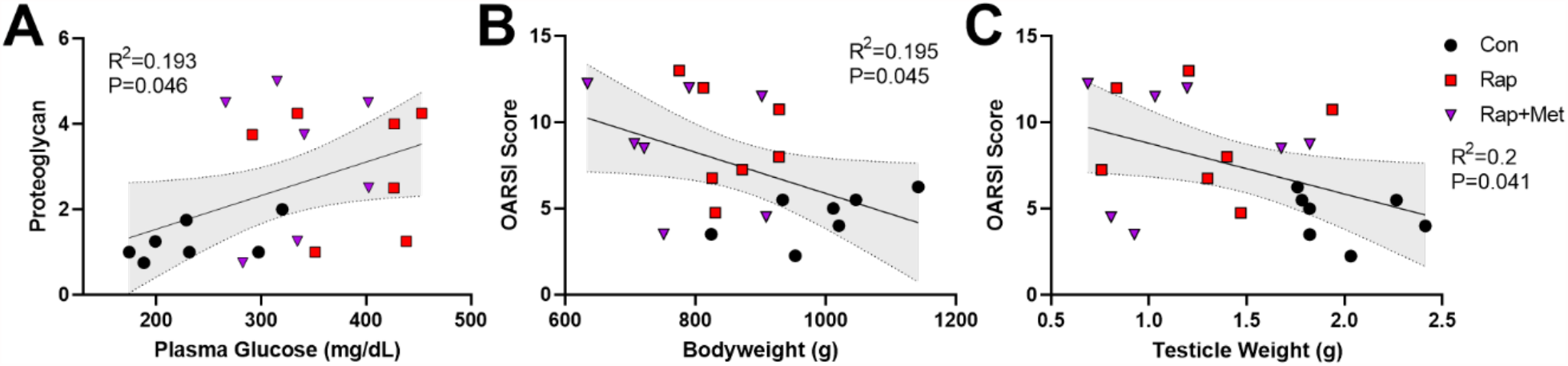
Proteoglycan loss correlated with hyperglycemia. Correlations between proteoglycan loss and plasma glucose (A), bodyweight and total OARSI score (B), and testicle weight and total OARSI score (C) are shown. Shaded bands represent 95% CI.

### Effects of rapamycin and rapamycin+metformin on mTOR, AMPK, and protease expression

To evaluate mTORC1 signaling in articular cartilage, we measured the phosphorylation of ribosomal protein S6 (P-RPS6) at Ser235/236 using IHC and western blotting. Representative images of P-RPS6 IHC are shown in Figure 4A. P-RPS6 was decreased by 90-95% in the medial tibial articular cartilage of Rap and Rap+Met treated guinea pigs as assessed by percentage of P-RPS6-positive cells (P=0.001 for Rap, P=0.01 for Rap+Met; Figure 4B), and by staining intensity (P=0.02 for both; Figure 4C). mTORC1 inhibition was further supported by an 81% lower ratio of phosphorylated to total RPS6 in glenohumeral cartilage from Rap (P=0.005; Figure 4E). Rap+Met trended to decrease RPS6 phosphorylation by 48% (P=0.06). There were no signficant effect on the phosphorylation of the mTORC2 substrate Akt at Ser473 in Rap or Rap+Met compared to control (Figure 4F; P=0.11). AMPK activity was measured using western blot to assess phosphorylation of AMPK at Thr172 (P-AMPK). P-AMPK was not changed by Rap alone (P=0.83; Figure 4G) but was elevated 77% by Rap+Met (P=0.05). Rap or Rap+Met did not significantly change the conversion of LC3B I to II (P>0.99 for both; Figure 4H) nor a disintegrin and metalloproteinase with thrombospondin motifs 5 (ADAMTS5; Figure 4I; P=0.97 for Rap, P=0.35 for Rap+Met). Matrix metalloproteinase 13 (MMP13) was unchanged by Rap (P>0.99) but trended higher in Rap+Met (P=0.09; Figure 4J).

**Figure 4:**
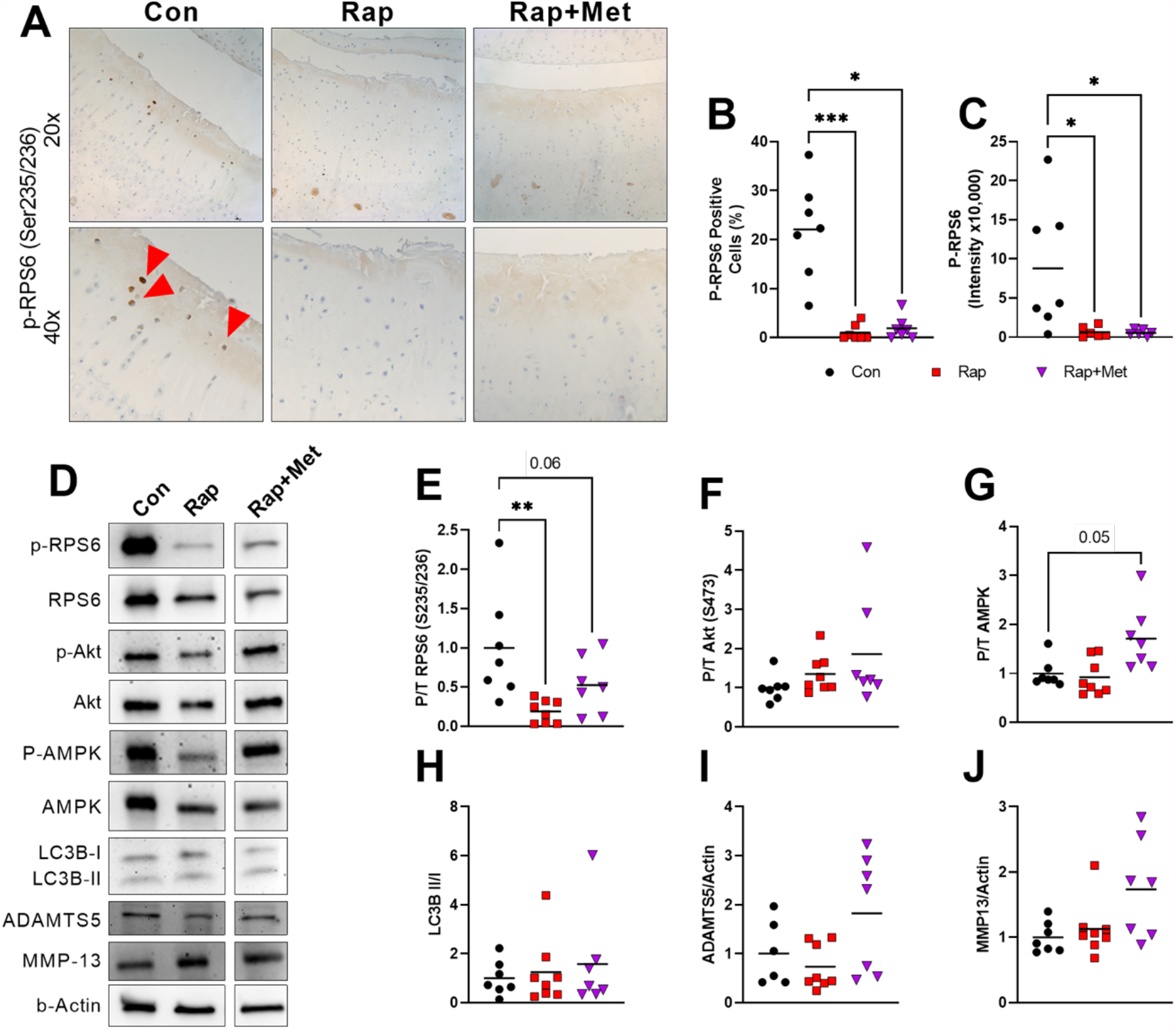
Rapamycin and rapamycin plus metformin inhibited mTORC1 but had no effect on mTORC2 or autophagy. IHC was performed on the medial tibia for P-RPS6 (A; n=7 per group) and quantified as percent positive cells (B) and mean integrated intensity (C). Red arrowheads indicate cells staining positive for P-RPS6. Western blot was performed on glenohumeral cartilage (D) for P-RPS6 (E), P-Akt (F), P-AMPK (G), LC3B (H), ADAMTS5 (I), and MMP-13 (J). n=8 per group for Rap and n=7 per group for Con and Rap+Met. Images are outlined in black to show that, while each band is from the same blot, bands were selected for presentation to best represent the mean change. *P<0.05 vs Con, **P<0.01 vs Con.

### Rapamycin and rapamycin+metformin decreased subchondral and diaphyseal bone thickness

Representative microCT images shown in Figure 5A were used to quantify the effect of experimental diets on subchondral bone parameters. Mean subchondral cortical thickness was decreased by Rap and Rap+Met in the medial (29%, P=0.003 for Rap; 23%, P=0.007 for Rap+Met) and lateral (21% for Rap; 20% for Rap+Met; P=0.01 for both) tibia (Figure 5B). Rap and Rap+Met decreased trabecular spacing by 15% and 16%, respectively, in the lateral tibia only (P=0.006 for both; Figure S2B). Trabecular thickness, trabecular spacing in other compartments, and bone volume fraction were not affected by any experimental diet (Figures S2A-C). Further investigation revealed that cortical thickness at the femoral diaphysis was decreased by Rap (P=0.001) and Rap+Met (P=0.01; Fig 5C), and this change was proportionate to the decrease observed in the medial tibial subchondral bone (Figure 5D). Further, medial tibial cortical thickness was correlated to bodyweight (R^2^=0.47, P=0.01; Figure 5E), suggesting the smaller body mass of Rap and Rap+Met treated guinea pigs may have contributed to decreased cortical thickness. Femoral epicondylar width (Figure 5F) was not statistically different between groups (Rap, P=0.42; Rap+Met, P=0.45), suggesting our treatments did not affect skeletal development.

**Figure 5:**
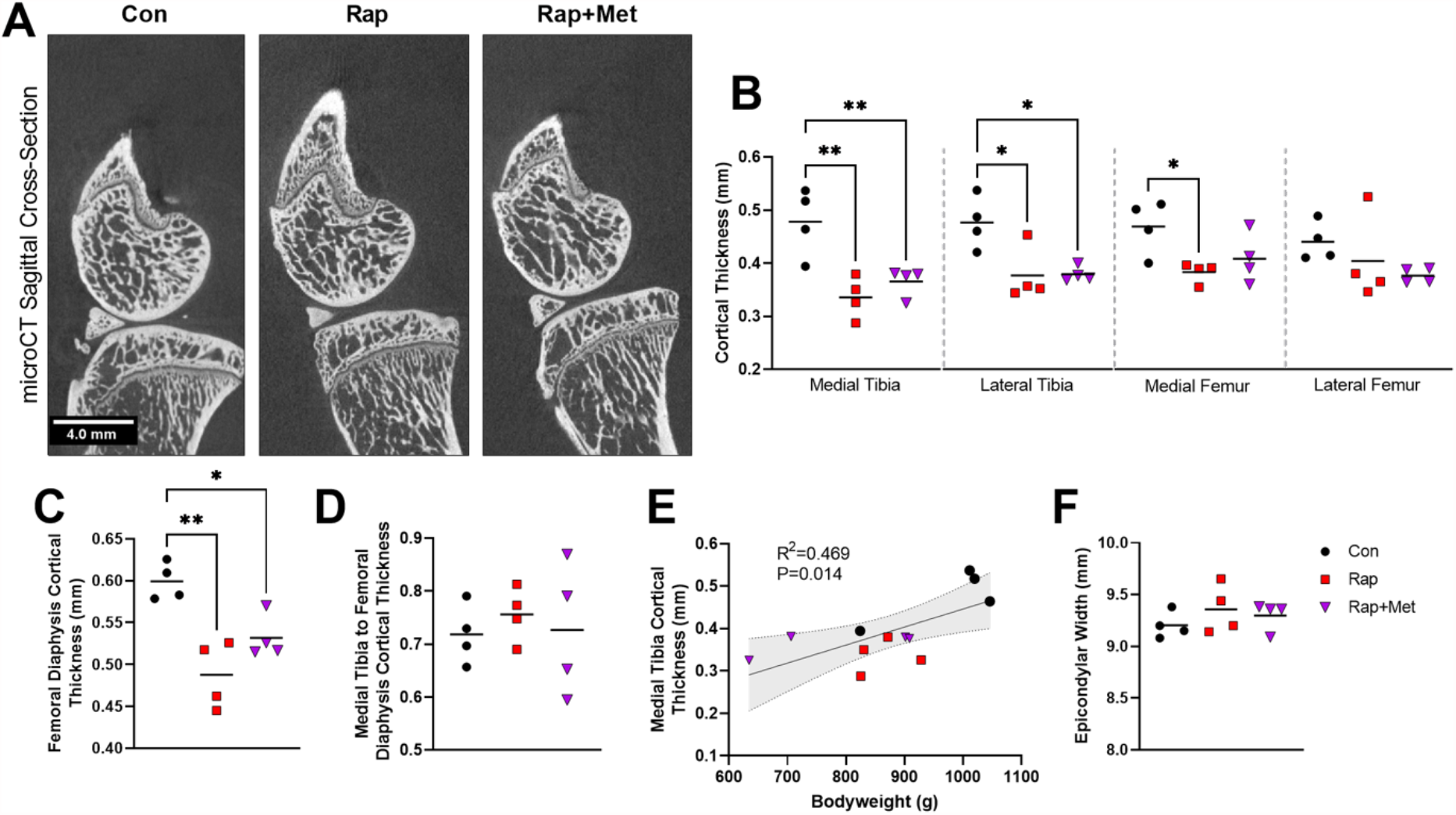
Decreased subchondral bone thickness in rapamycin and rapamycin plus metformin treated guinea pigs. Representative microCT sagittal cross sections from the medial aspect of the joint are shown (A). Subchondral cortical thickness was measured in the medial and lateral tibial plateaus and femoral condyles (B), and cortical thickness was measured in the femoral diaphysis (C). Medial tibial cortical thickness relative to femoral diaphyseal cortical thickness was found to be similar between groups (D). Medial tibial cortical thickness was highly correlated to bodyweight (E). Femoral epicondylar width was found to be similar between groups (F). N=4 per group. Shaded bands represent 95% CI. *P<0.05 vs Con, **P<0.01 vs Con.

## Discussion

The purpose of this study was to test if dietary Rap or Rap+Met could delay the onset of age-related OA in the outbred Dunkin-Hartley guinea pig. We found that at concentrations shown to extend lifespan, dietary Rap and Rap+Met inhibited mTORC1 but not mTORC2 signaling in articular cartilage, and Rap+Met increased AMPK phosphorylation. Surprisingly, guinea pigs treated with Rap, with or without Met, developed greater age-related OA compared to control. Guinea pigs receiving Rap and Rap+Met also displayed increased plasma glucose, which correlated with proteoglycan loss. These findings indicate that off-target side effects of Rap are associated with greater OA pathology. Further, in the face of these Rap-induced side effects, mTORC1 inhibition may not slow the progression of age-related OA in Dunkin Harltey guinea pigs.

Despite inhibiting mTORC1 in articular cartilage, our findings indicate that guinea pigs treated with Rap, with or without Met, had exacerbated age-related OA in the medial tibial plateau. Further, Rap and Rap+Met treated guinea pigs had greater total OARSI scores even though they weighed less, which is contrary to previous work where lower body weight was accompanied by lower OA scores in guinea pigs (29). Although there is precedent that mTORC1 inhibition by intra-articular injection of Rap is associated with exacerbated temporomandibular joint (TMJ) OA (44), our findings were opposite of our original hypothesis and previous results using Rap in secondary models of knee OA (18,19). The guinea pigs in the current study received a dose of Rap that achieved similar circulating Rap concentrations shown to extend lifespan in mice (14). Additionally, the dose of Rap in guinea pigs was similar to the dose shown to protect against secondary OA in mice (0.7 vs 1 mg/kg/day in guinea pigs vs. mice) (18). These findings suggest that dose of Rap was not a likely factor contributing to differences between studies. In our study, Rap and Rap+Met treatment inhibited mTORC1 but not mTORC2 in articular cartilage. Previous work has shown that deleting articular cartilage mTOR (21) or treating with Rap (18,19) or Torin-1 (20) can attenuate secondary OA in mice and rabbits. These non-selective genetic and pharmacological methods likely disrupt the entire mTOR kinase and therefore could inhibit both mTORC1 and mTORC2 signaling. However, this remains speculative as mTORC2 signaling was not evaluated in these previous studies, and it continues to be unknown if mTORC2 inhibition is necessary for protection against either primary or secondary OA. In support of the notion that targeting mTORC2 modifies OA, inhibition of the mTORC2 substrate Akt protects against PTEN-deletion-induced OA by decreasing cellular senescence and oxidative stress (45). Further investigation is needed to resolve the role of each mTOR complex in the initiation, progression, and treatment of both primary and secondary OA.

Despite its lifespan-extending effects, chronic Rap treatment is commonly associated with several metabolic and immunological side effects including glucose intolerance, insulin resistance, hypertriglyceridemia, immunosuppression, testicular atrophy, lower body weight, and cataracts (17,39,46). Consistent with this, we showed that 12-weeks of dietary Rap and Rap+Met was accompanied by increased plasma glucose, testicular atrophy, and lower body weight. Despite increasing AMPK activity in articular cartilage and partially restoring glucose levels compared to Rap alone, the addition of Met to Rap did not offer protection against the detrimental effects of dietary Rap on OA pathology. The glucose lowering effects of Met are in line with previous studies where Met alleviated Rap-induced glucose intolerance only in female mice (37). However, our OA pathology findings are in contrast to previous studies that showed Met attenuated hyperglycemia-induced OA in mice (55). In our study, medial tibial proteoglycan loss was correlated with plasma glucose, and we propose that Rap-induced hyperglycemia may have contributed to worsened OA following dietary Rap treatment. In support of this hypothesis, diabetic mice show accelerated OA after injury, and chondrocytes cultured in high glucose media display decreased expression of Collagen II and increased MMP13 and inflammatory mediators IL-6 and NFkB (47,48). However, intermittent intraperitoneal injections of Rap lowered glucose and mitigated diabetes accelerated secondary OA (49). It is possible that Rap did offer partial protection against hyperglycemic stress but still resulted in greater OA pathology than control, as was observed by Ribeiro et al. (50). However, this remains speculative as we did not have a group exposed to hyperglycemic stress alone. Previous work suggests Rap can have divergent effects where it is beneficial in some diabetic models but causes adverse side effects in metabolically healthy models (17,51). Collectively, these data indicate that the adverse metabolic side-effects of dietary Rap treatment are associated with a deleterious impact on primary OA pathology and could limit the utility of systemic Rap as a healthspan extending treatment.

Rap has been implicated in attenuating secondary OA by increasing autophagy and decreasing protease expression (18,19). While autophagy is a highly dynamic process, the static marker of autophagy, LC3B, is commonly used as a surrogate for autophagic flux. In our study, we saw no effect by any treatment on LC3B or ADAMTS5, while Rap+Met trended to increase MMP13 in glenohumeral cartilage. Therefore, the inability to increase markers of autophagy and decrease proteases may be one contributing factor to why our lifespan-extending treatments did not protect and even worsened OA during aging and hyperglycemia. However, because proteoglycan loss was observed independent of increased protease expression in Rap-treated guinea pigs, decreased extracellular matrix (ECM) protein synthesis may have contributed to proteoglycan loss. More work is needed to determine the molecular and cellular mechanisms responsible for the deleterious effects of Rap and Rap+Met.

Treatment with Rap and Rap+Met also decreased subchondral cortical bone thickness in the medial and lateral tibia and the femoral diaphysis. As bone growth in guinea pigs ceases by 4 months (52), and epicondylar width was not different between groups, the differences in bone thickness were likely not the result of disrupted development. Decreased subchondral thickness was only observed in the tibia. Intra-articular injection of Rap into the TMJ caused subchondral bone loss by inhibiting pre-osteoblast proliferation (44), and Rap treatment also decreased osteoblast differentiation and bone matrix synthesis (53), which supports the idea that Rap can act directly on the bone to decrease thickness. However, we also found that subchondral thickness was highly correlated to bodyweight. This is in line with Wolff’s law and agrees with previous findings where bodyweight restriction decreased cortical bone thickness in the femoral diaphysis (54). Therefore, both local and systemic effects of Rap likely contributed to reduced cortical bone thickness.

Although we provide new insight into the role of mTOR during primary OA progression, we recognize some study limitations. While the guinea pig is an excellent model of primary OA, it is not a widespread model for biomedical research and molecular probes are seldom designed for reactivity with guinea pig tissue. Due to reactivity issues with IHC in guinea pig cartilage (Figure S3), some of our analyses relied on western blot from glenohumeral cartilage. Although guinea pigs also develop mild glenohumeral OA(30), this is not the site at which we measured OA pathology. Our study could not conclusively determine if the deleterious effects of Rap stemmed from its direct effects on the joint or off-target effects on other tissues. However, our data suggest hyperglycemia induced by off-target actions of Rap was associated with worsened age-related OA. The Dunkin Hartley guinea pig is an outbred model of primary OA which leads to inherent variability. While this could be perceived as a limitation, we contend that the variability and the choice of animal model adds translational value since this more closely recapitulates the genetic diversity and OA heterogeneity in humans. We acknowledge that although the sample size used in our study was in line with previous studies using guinea pigs, the varability could have possibly limited our ability to detect more subtle differences between groups. However, this does not detract from the findings that guinea pigs treated with both Rap and Rap+Met had worse OA. Further, the presence of largely overlapping and consistent deleterious outcomes in both groups receiving Rap increases our confidence that the side effects accompanying Rap contribute to worsened primary OA.

## Conclusion

In summary, we have shown that at doses previously shown to extend lifespan, dietary Rap and Rap+Met caused hyperglycemia and was associated with aggravated OA in Dunkin Hartley guinea pigs despite inhibiting mTORC1 in articular cartilage. Treatments that extend lifespan without a proportional delay in age-related chronic diseases and disabilities is counter to the concept of healthspan extension. Our findings that guinea pigs treated with Rap had worse OA pathology raises concerns regarding the efficacy of dietary Rap as a life- and healthspan-extending agent. Additional work is needed to investigate the role of alternative routes of administration or Rap anaologs that may capture the positive benefits of Rap while minimizing off-target effects. Our data also reveal that the contribution of mTOR in articular cartilage and chondrocyte metabolism is incompletely understood and additional research is needed to clarify the individual and combined role of mTORC1 and mTORC2 signaling in both primary and secondary OA.

## Abbreviations

OA: osteoarthritis
mTOR: mechanistic target of rapamycin
AMPK: AMP-activated protein kinase
Rap: rapamycin
Met: metformin
mTORC1: mTOR complex I
mTORC2: mTOR complex II
NBF: neutral buffered formalin µ
CT: micro computed tomography
ROI: region of interest
IHC: immunohistochemistry
OARSI: osteoarthritis research society international
SD: standard deviation
RPS6: ribosomal protein S6
ADAMTS5: a disintegrin and metalloproteinase with thrombospondin motifs 5
MMP13: matrix metalloproteinase 13
TMJ: temporomandibular joint
IL-6: interleukin 6
NFkB: nuclear factor kappa-light-chain-enhancer of activated B-cells
ECM: extracellular matrix

## Acknowledgements

The authors would like to thank Greg Friesenhahn at the Analytical Pharmacology and Drug Evaluation Core of the San Antonio Nathan Shock Center. We would also like to acknowledge the technical assistance from William Fairfield, Oscar Safairad, Alex Nichol, Nathan Carper, and Morgan Berland. We also acknowledge the assistance of the University of Wisconsin Translational Research Initiatives in Pathology (TRIP) laboratory supported by the UW Department of Pathology and Laboratory Medicine, UWCCC (P30 CA014520).

## Author Contributions

Study design: AK. Data collection: DM, CE, AK, KS, MJ. Data analysis and interpretation: DM, AK. Manuscript preparation: DM, AK. All authors approved of the final manuscript.

## Availabililty of Data and Materials

Data from this study are available from the corresponding author upon reasonable request.

## Ethics Approval

Animal use was approved by the University of Illinois at Urbana-Champaign IRB and IACUC.

## Consent for Publication

Not applicable.

## Supplementary Material

### Supplementary Figure Legends

**Figure S1:**
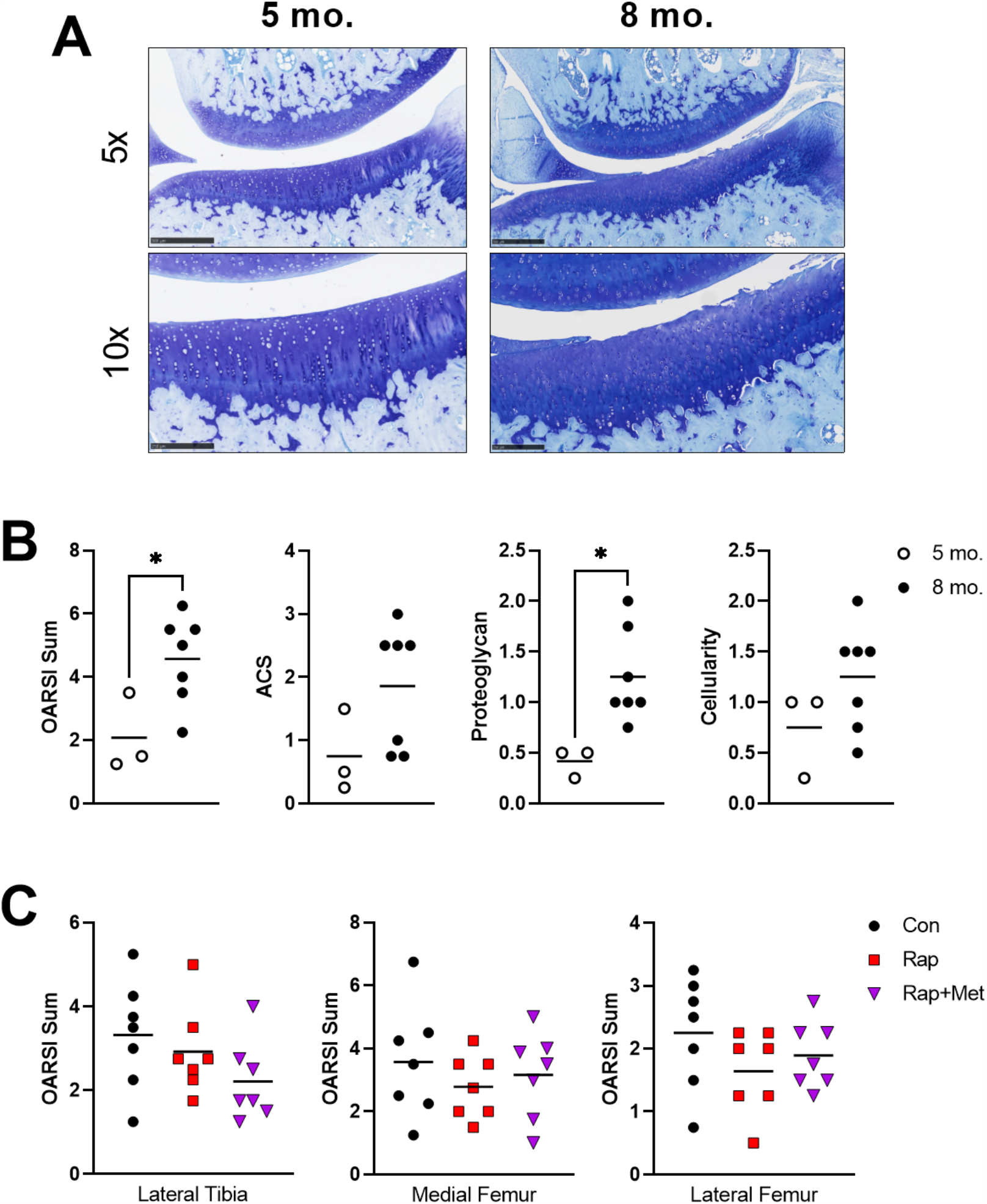
OA pathology increased from 5- to 8-months of age. Total OARSI scores are shown from the lateral tibia, medial femur, and lateral femur (A). Histological images of knee joints from 5- and 8-month-old guinea pigs (B; scale bars are 0.5mm and 0.25mm in 5x and 10x images, respectively) were graded for total OARSI score and individual OARSI criteria (C). N=3 for 5-month and N=7 for 8-month. *P<0.05 vs Con.

**Figure S2:**
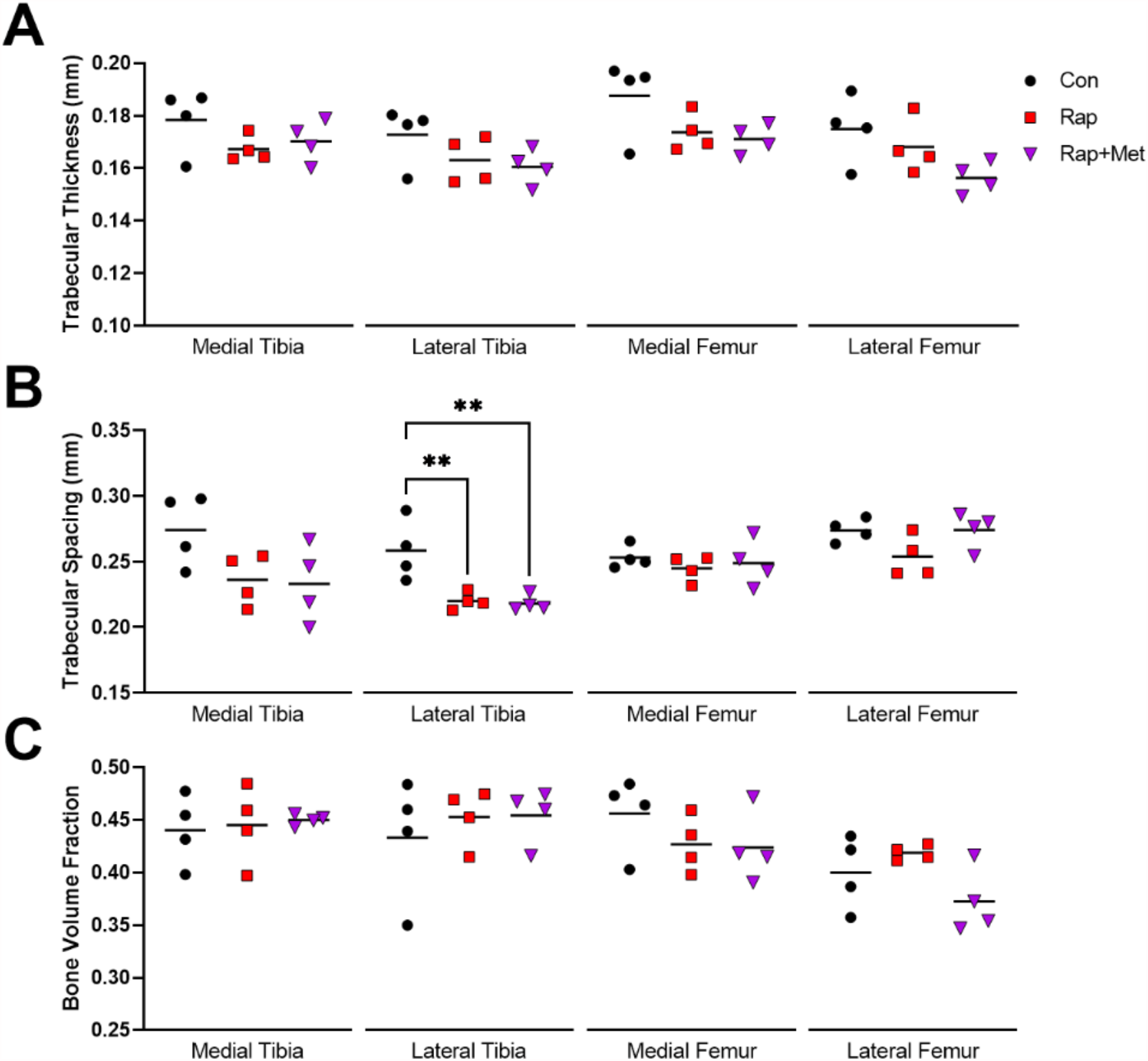
Trabecular bone changes in response to experimental diets. Trabecular thickness (A), spacing (B), and bone volume fraction (C) were measured using microCT. N=4 per group. *P<0.05 vs Con.

**Figure S3:**
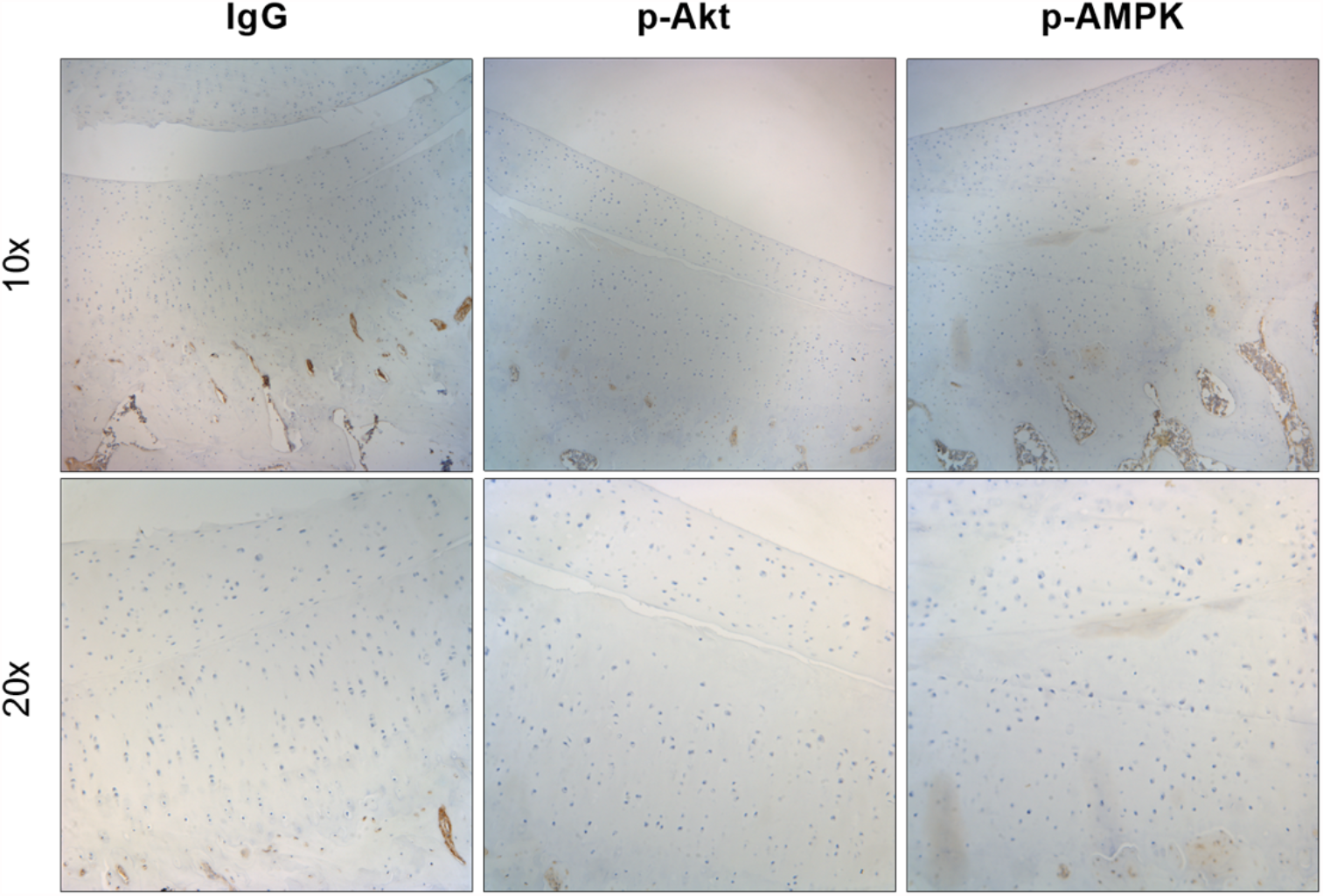
Antibody reactivity with guinea pig articular cartilage was limited. Immunohistochemical staining was performed, and no reactivity was observed using primary antibodies against P-Akt Ser473 or P-AMPK Thr172.

## References

1. Brown TD, Johnston RC, Saltzman CL, Marsh JL, Buckwalter JA. Posttraumatic Osteoarthritis: A First Estimate of Incidence, Prevalence, and Burden of Disease. J Orthop Trauma. 2006 Nov;20(10):739–44. Available from: https://pubmed.ncbi.nlm.nih.gov/17106388/

2. Aki T, Hashimoto K, Ogasawara M, Itoi E. A whole-genome transcriptome analysis of articular chondrocytes in secondary osteoarthritis of the hip. Agarwal S, editor. PLoS One. 2018 Jun 26;13(6):e0199734. Available from: https://pubmed.ncbi.nlm.nih.gov/29944724/

3. Xu Y, Barter MJ, Swan DC, Rankin KS, Rowan AD, Santibanez-Koref M, et al. Identification of the pathogenic pathways in osteoarthritic hip cartilage: commonality and discord between hip and knee OA. Osteoarthr Cartil. 2012 Sep;20(9):1029–38. Available from: https://pubmed.ncbi.nlm.nih.gov/22659600/

4. Moon PM, Shao ZY, Wambiekele G, Appleton CTG, Laird DW, Penuela S, et al. Global Deletion of Pannexin 3 Resulting in Accelerated Development of Aging-Induced Osteoarthritis in Mice. Arthritis Rheumatol. 2021 May 25; Available from: https://pubmed.ncbi.nlm.nih.gov/33426805/

5. Yu D, Hu J, Sheng Z, Fu G, Wang Y, Chen Y, et al. Dual roles of misshapen/NIK-related kinase (MINK1) in osteoarthritis subtypes through the activation of TGFβ signaling. Osteoarthr Cartil. 2020 Jan;28(1):112–21. Available from: https://pubmed.ncbi.nlm.nih.gov/31647983/

6. Bouderlique T, Vuppalapati KK, Newton PT, Li L, Barenius B, Chagin AS. Targeted deletion of Atg5 in chondrocytes promotes age-related osteoarthritis. Ann Rheum Dis. 2016 Mar;75(3):627–31. Available from: https://pubmed.ncbi.nlm.nih.gov/26438374/

7. O’Conor CJ, Ramalingam S, Zelenski NA, Benefield HC, Rigo I, Little D, et al. Cartilage-specific knockout of the mechanosensory ion channel TRPV4 decreases age-related osteoarthritis. Sci Rep. 2016;6(July):1–10. Available from: http://dx.doi.org/10.1038/srep29053

8. Usmani SE, Ulici V, Pest MA, Hill TL, Welch ID, Beier F. Context-specific protection of TGFα null mice from osteoarthritis. Sci Rep. 2016 Sep 26;6(1):1–11. Available from: https://pubmed.ncbi.nlm.nih.gov/27457421/

9. Loeser RF, Kelley KL, Armstrong A, Collins JA, Diekman BO, Carlson CS. Deletion of JNK Enhances Senescence in Joint Tissues and Increases the Severity of Age-Related Osteoarthritis in Mice. Arthritis Rheumatol. 2020 Oct 26;72(10):1679–88. Available from: https://pubmed.ncbi.nlm.nih.gov/32418287/

10. Zhang H, Wang H, Zeng C, Yan B, Ouyang J, Liu X, et al. mTORC1 activation downregulates FGFR3 and PTH/PTHrP receptor in articular chondrocytes to initiate osteoarthritis. Osteoarthr Cartil. 2017 Jun;25(6):952–63. Available from: https://pubmed.ncbi.nlm.nih.gov/28043938/

11. Zhou S, Lu W, Chen L, Ge Q, Chen D, Xu Z, et al. AMPK deficiency in chondrocytes accelerated the progression of instability-induced and ageing-associated osteoarthritis in adult mice. Sci Rep. 2017 Apr 22;7(1):43245. Available from: https://pubmed.ncbi.nlm.nih.gov/28225087/

12. Johnson SC, Rabinovich PS, Kaeberlin M. mTOR is a key modulator of ageing and age-related disease. Nature. 2013;493(7432):338–45. Available from: https://pubmed.ncbi.nlm.nih.gov/23325216/

13. Salminen A, Kaarniranta K, Kauppinen A. Age-related changes in AMPK activation: Role for AMPK phosphatases and inhibitory phosphorylation by upstream signaling pathways. Ageing Res Rev. 2016;28:15–26. Available from: https://pubmed.ncbi.nlm.nih.gov/27060201/

14. Harrison DE, Strong R, Sharp ZD, Nelson JF, Astle CM, Flurkey K, et al. Rapamycin fed late in life extends lifespan in genetically heterogeneous mice. Nature. 2009;460(7253):392–5. Available from: https://pubmed.ncbi.nlm.nih.gov/19587680/

15. Strong R, Miller RA, Antebi A, Astle CM, Bogue M, Denzel MS, et al. Longer lifespan in male mice treated with a weakly estrogenic agonist, an antioxidant, an α-glucosidase inhibitor or a Nrf2-inducer. Aging Cell. 2016 Oct 16;15(5):872–84. Available from: https://pubmed.ncbi.nlm.nih.gov/27312235/

16. Barzilai N, Crandall JP, Kritchevsky SB, Espeland MA. Metformin as a Tool to Target Aging. Cell Metab. 2016;23(6):1060–5. Available from: http://dx.doi.org/10.1016/j.cmet.2016.05.011

17. Lamming DW, Ye L, Katajisto P, Goncalves MD, Saitoh M, Stevens DM, et al. Rapamycin-Induced Insulin Resistance Is Mediated by mTORC2 Loss and Uncoupled from Longevity. Science (80-). 2012 Mar 30;335(6076):1638–43. Available from: https://pubmed.ncbi.nlm.nih.gov/22461615/

18. Caramés B, Hasegawa A, Taniguchi N, Miyaki S, Blanco FJ, Lotz M. Autophagy activation by rapamycin reduces severity of experimental osteoarthritis. Ann Rheum Dis. 2012 Apr 4;71(4):575–81. Available from: https://pubmed.ncbi.nlm.nih.gov/22084394/

19. Takayama K, Kawakami Y, Kobayashi M, Greco N, Cummins JH, Matsushita T, et al. Local intra-articular injection of rapamycin delays articular cartilage degeneration in a murine model of osteoarthritis. Arthritis Res Ther. 2014 Dec 17;16(6):482. Available from: https://pubmed.ncbi.nlm.nih.gov/25403236/

20. Cheng N-T, Guo A, Cui Y-P. Intra-articular injection of Torin 1 reduces degeneration of articular cartilage in a rabbit osteoarthritis model. Bone Joint Res. 2016 Jun;5(6):218–24. Available from: https://pubmed.ncbi.nlm.nih.gov/27301478/

21. Zhang Y, Vasheghani F, Li Y, Blati M, Simeone K, Fahmi H, et al. Cartilage-specific deletion of mTOR upregulates autophagy and protects mice from osteoarthritis. Ann Rheum Dis. 2015 Jul;74(7):1432–40. Available from: https://pubmed.ncbi.nlm.nih.gov/24651621/

22. Chen J, Crawford R, Xiao Y. Vertical inhibition of the PI3K/Akt/mTOR pathway for the treatment of osteoarthritis. J Cell Biochem. 2013;114(2):245–9.

23. Pal B, Endisha H, Zhang Y, Kapoor M. mTOR: A potential therapeutic target in osteoarthritis? Drugs R D. 2015;15(1):27–36. Available from: https://pubmed.ncbi.nlm.nih.gov/25688060/

24. Chen K, Lin ZW, He S mao, Wang C qiang, Yang J cheng, Lu Y, et al. Metformin inhibits the proliferation of rheumatoid arthritis fibroblast-like synoviocytes through IGF-IR/PI3K/AKT/m-TOR pathway. Biomed Pharmacother. 2019;115(April 2018):1–8. Available from: https://pubmed.ncbi.nlm.nih.gov/31028998/

25. Wang C, Yang Y, Zhang Y, Liu J, Yao Z, Zhang C. Protective effects of metformin against osteoarthritis through upregulation of SIRT3-mediated PINK1/Parkin-dependent mitophagy in primary chondrocytes. Biosci Trends. 2018 Dec 31;12(6):605–12. Available from: https://pubmed.ncbi.nlm.nih.gov/30584213/

26. Petursson F, Husa M, June R, Lotz M, Terkeltaub R, Liu-Bryan R. Linked decreases in liver kinase B1 and AMP-activated protein kinase activity modulate matrix catabolic responses to biomechanical injury in chondrocytes. Arthritis Res Ther. 2013;15(4):R77. Available from: https://pubmed.ncbi.nlm.nih.gov/23883619/

27. Li J, Zhang B, Liu W-X, Lu K, Pan H, Wang T, et al. Metformin limits osteoarthritis development and progression through activation of AMPK signalling. Ann Rheum Dis. 2020 May;79(5):635–45. Available from: https://pubmed.ncbi.nlm.nih.gov/32156705/

28. Wang Y, Hussain SM, Wluka AE, Lim YZ, Abram F, Pelletier J-P, et al. Association between metformin use and disease progression in obese people with knee osteoarthritis: data from the Osteoarthritis Initiative—a prospective cohort study. Arthritis Res Ther. 2019 Dec 24;21(1):127. Available from: https://pubmed.ncbi.nlm.nih.gov/31126352/

29. Bendele AM, Hulman JF. Effects of Body Weight Restriction on the Development and Progression of Spontaneous Osteoarthritis in Guinea Pigs. Arthritis Rheum. 1991 Oct 7;34(9):1180–4. Available from: https://pubmed.ncbi.nlm.nih.gov/1930336/

30. Bendele AM, White SL, Hulman JF. Osteoarthrosis in guinea pigs: histopathologic and scanning electron microscopic features. Lab Anim Sci. 1989;39(2):115–21. Available from: https://pubmed.ncbi.nlm.nih.gov/2709799/

31. Radakovich LB, Marolf AJ, Shannon JP, Pannone SC, Sherk VD, Santangelo KS. Development of a microcomputed tomography scoring system to characterize disease progression in the Hartley guinea pig model of spontaneous osteoarthritis. Connect Tissue Res. 2018 Nov 2;59(6):523–33. Available from: https://pubmed.ncbi.nlm.nih.gov/29226725/

32. Kraus VB, Huebner JL, DeGroot J, Bendele A. The OARSI histopathology initiative - recommendations for histological assessments of osteoarthritis in the guinea pig. Osteoarthr Cartil. 2010;18(SUPPL. 3):S35–52. Available from: http://dx.doi.org/10.1016/j.joca.2010.04.015

33. Bendele AM. Animal models of osteoarthritis. In: Journal of Musculokeletal Neuron Interaction. 2001. p. 363–76. Available from: https://pubmed.ncbi.nlm.nih.gov/15758487/

34. Radakovich LB, Marolf AJ, Culver LA, Santangelo KS. Calorie restriction with regular chow, but not a high-fat diet, delays onset of spontaneous osteoarthritis in the Hartley guinea pig model. Arthritis Res Ther. 2019 Dec 13;21(1):145. Available from: https://pubmed.ncbi.nlm.nih.gov/31196172/

35. Martin-Montalvo A, Mercken EM, Mitchell SJ, Palacios HH, Mote PL, Scheibye-Knudsen M, et al. Metformin improves healthspan and lifespan in mice. Nat Commun. 2013 Oct 16;4(1):2192. Available from: https://pubmed.ncbi.nlm.nih.gov/23900241/

36. Miller RA, Harrison DE, Astle CM, Baur JA, Boyd AR, de Cabo R, et al. Rapamycin, But Not Resveratrol or Simvastatin, Extends Life Span of Genetically Heterogeneous Mice. Journals Gerontol Ser A. 2011 Feb;66A(2):191–201. Available from: https://pubmed.ncbi.nlm.nih.gov/20974732/

37. Weiss R, Fernandez E, Liu Y, Strong R, Salmon AB. Metformin reduces glucose intolerance caused by rapamycin treatment in genetically heterogeneous female mice. Aging (Albany NY). 2018 Mar 22;10(3):386–401. Available from: https://pubmed.ncbi.nlm.nih.gov/29579736/

38. De Oliveira MA, Martins E Martins F, Wang Q, Sonis S, Demetri G, George S, et al. Clinical presentation and management of mTOR inhibitor-associated stomatitis. Oral Oncol. 2011;47(10):998–1003. Available from: https://pubmed.ncbi.nlm.nih.gov/21890398/

39. Wilkinson JE, Burmeister L, Brooks S V., Chan C-C, Friedline S, Harrison DE, et al. Rapamycin slows aging in mice. Aging Cell. 2012 Aug;11(4):675–82. Available from: http://doi.wiley.com/10.1111/j.1474-9726.2012.00832.x

40. Fraser KW. Effect of storage in formalin on organ weights of rabbits. New Zeal J Zool. 1985;12(2):169–74.

41. Tardif S, Ross C, Bergman P, Fernandez E, Javors M, Salmon A, et al. Testing Efficacy of Administration of the Antiaging Drug Rapamycin in a Nonhuman Primate, the Common Marmoset. Journals Gerontol Ser A Biol Sci Med Sci. 2015 May;70(5):577–88. Available from: https://pubmed.ncbi.nlm.nih.gov/25038772/

42. Fernandez E, Ross C, Liang H, Javors M, Tardif S, Salmon AB. Evaluation of the pharmacokinetics of metformin and acarbose in the common marmoset. Pathobiol Aging Age-related Dis. 2019;9(1):1657756. Available from: https://pubmed.ncbi.nlm.nih.gov/31497263/

43. Buie HR, Campbell GM, Klinck RJ, MacNeil JA, Boyd SK. Automatic segmentation of cortical and trabecular compartments based on a dual threshold technique for in vivo micro-CT bone analysis. Bone. 2007 Oct;41(4):505–15. Available from: https://pubmed.ncbi.nlm.nih.gov/17693147/

44. Li Y, Yang J, Liu Y, Yan X, Zhang Q, Chen J, et al. Inhibition of mTORC1 in the rat condyle subchondral bone aggravates osteoarthritis induced by the overly forward extension of the mandible. Am J Transl Res. 2021;13(1):270–85. Available from: https://pubmed.ncbi.nlm.nih.gov/33527023/

45. Xie J, Lin J, Wei M, Teng Y, He Q, Yang G, et al. Sustained Akt signaling in articular chondrocytes causes osteoarthritis via oxidative stress-induced senescence in mice. Bone Res. 2019 Dec 5;7(1):23. Available from: https://pubmed.ncbi.nlm.nih.gov/31646013/

46. Aggarwal D, Fernandez ML, Soliman GA. Rapamycin, an mTOR inhibitor, disrupts triglyceride metabolism in guinea pigs. Metabolism. 2006 Jun;55(6):794–802. Available from: https://pubmed.ncbi.nlm.nih.gov/16713440/

47. Chen YJ, Chan DC, Lan KC, Wang CC, Chen CM, Chao SC, et al. PPARγ is involved in the hyperglycemia-induced inflammatory responses and collagen degradation in human chondrocytes and diabetic mouse cartilages. J Orthop Res. 2015;33(3):373–81. Available from: https://pubmed.ncbi.nlm.nih.gov/25410618/

48. Liang H, Wang H, Luo L, Fan S, Zhou L, Liu Z, et al. Toll-like receptor 4 promotes high glucose-induced catabolic and inflammatory responses in chondrocytes in an NF-κB-dependent manner. Life Sci. 2019;228(April):258–65. Available from: https://pubmed.ncbi.nlm.nih.gov/30953645/

49. Ribeiro M, López de Figueroa P, Nogueira-Recalde U, Centeno A, Mendes AF, Blanco FJ, et al. Diabetes-accelerated experimental osteoarthritis is prevented by autophagy activation. Osteoarthr Cartil. 2016;24(12):2116–25.

50. Ribeiro M, López de Figueroa P, Nogueira-Recalde U, Centeno A, Mendes AF, Blanco FJ, et al. Diabetes-accelerated experimental osteoarthritis is prevented by autophagy activation. Osteoarthr Cartil. 2016;24(12):2116–25. Available from: https://pubmed.ncbi.nlm.nih.gov/27390029/

51. Reifsnyder PC, Flurkey K, Te A, Harrison DE. Rapamycin treatment benefits glucose metabolism in mouse models of type 2 diabetes. Aging (Albany NY). 2016;8(11):3120–30. Available from: https://pubmed.ncbi.nlm.nih.gov/27922820/

52. Watson PJ, Hall LD, Malcolm A, Tyler JA. Degenerative joint disease in the guinea pig: Use of magnetic resonance imaging to monitor progression of bone pathology. Arthritis Rheum. 1996;39(8):1327–37. Available from: https://pubmed.ncbi.nlm.nih.gov/8702441/

53. Xian L, Wu X, Pang L, Lou M, Rosen C, Qui T, et al. Matrix IGF-1 regulates bone mass by activation of mTOR in mesenchymal stem cells. Nat Med. 2012;18(7):1095–101. Available from: https://pubmed.ncbi.nlm.nih.gov/22729283/

54. Hamrick MW, Ding KH, Ponnala S, Ferrari SL, Isales CM. Caloric restriction decreases cortical bone mass but spares trabecular bone in the mouse skeleton: Implications for the regulation of bone mass by body weight. J Bone Miner Res. 2008;23(6):870–8. Available from: https://pubmed.ncbi.nlm.nih.gov/18435579/

55. Dawood AF, Alzamil N, Ebrahim HA, Abdel Kader DH, Kamar SS, Haidara MA, et al. Metformin pretreatment suppresses alterations to the articular cartilage ultrastructure and knee joint tissue damage secondary to type 2 diabetes mellitus in rats. Ultrastruct Pathol. 2020;44(3):273–82. Available from: https://pubmed.ncbi.nlm.nih.gov/32404018/

56. Choi YH, Sang KG, Lee MG. Dose-independent pharmacokinetics of metformin in rats: Hepatic and gastrointestinal first-pass effects. J Pharm Sci. 2006;95(11):2543–52. Available from: https://pubmed.ncbi.nlm.nih.gov/16937336/

57. Graham GG, Punt J, Arora M, Day RO, Doogue MP, Duong JK, et al. Clinical pharmacokinetics of metformin. Clin Pharmacokinet. 2011;50(2):81–98. Available from: https://pubmed.ncbi.nlm.nih.gov/21241070/

